# Multiscale analysis and optimal glioma therapeutic candidate discovery using the CANDO platform

**DOI:** 10.1101/2025.05.19.654757

**Authors:** Sumei Xu, William Mangione, Melissa Van Norden, Katherine Elefteriou, Yakun Hu, Zackary Falls, Ram Samudrala

## Abstract

Glioma is a highly malignant brain tumor with limited treatment options. We employed the Computational Analysis of Novel Drug Opportunities (CANDO) platform for multiscale therapeutic discovery to predict new glioma therapies. We began by computing interaction scores between extensive libraries of drugs/compounds and proteins to generate “interaction signatures” that model compound behavior on a proteomic scale. Compounds with signatures most similar to those of drugs approved for a given indication were considered potential treatments. These compounds were further ranked by degree of consensus in corresponding similarity lists. We benchmarked performance by measuring the recovery of approved drugs in these similarity and consensus lists at various cutoffs, using multiple metrics and comparing results to random controls and performance across all indications. Compounds ranked highly by consensus but not previously associated with the indication of interest were considered new predictions. Our benchmarking results showed that CANDO improved accuracy in identifying glioma-associated drugs across all cutoffs compared to random controls. Our predictions, supported by literature-based analysis, identified 23 potential glioma treatments, including approved drugs like vitamin D, taxanes, vinca alkaloids, topoisomerase inhibitors, and folic acid, as well as investigational compounds such as ginsenosides, chrysin, resiniferatoxin, and cryptotanshinone. Further functional annotation-based analysis of the top targets with the strongest interactions to these predictions identified Vitamin D3 receptor, thyroid hormone receptor, acetylcholinesterase, cyclin-dependent kinase 2, tubulin alpha chain, dihydrofolate reductase, and thymidylate synthase. These findings indicate that CANDO’s multitarget, multiscale framework is effective in identifying glioma drug candidates thereby informing new strategies for improving treatment.

## 1 Introduction

Glioma is one of the most aggressive and fatal forms of malignant brain tumors, particularly prevalent among the elderly, with high rates of occurrence and mortality [1, 2]. Currently, chemotherapy is the primary treatment for glioma due to its aggressive progression, various pathologies, and the challenges associated with complete surgical removal [3, 4]. However, the effectiveness of chemotherapy is significantly limited by factors such as the selective permeability of the blood-brain barrier (BBB), neurotoxicity, and inadequate drug delivery to the tumor site [5–9]. Furthermore, the ATP-dependent efflux transporter, P-glycoprotein (P-gp), located on the BBB, contributes to the removal of chemotherapeutic agents [10]. A substantial proportion of patients with glioma (about 90%) experience tumor recurrence in the local area after initial treatment [11]. Unfortunately, effective therapeutic options for recurrent glioma are lacking. As a result, there is an urgent need to advance our understanding of the molecular pathology of glioma, identify new therapeutic targets, and develop innovative treatment strategies. A major challenge in modern medicine is the limited availability of new glioma drugs that can cross the BBB [12–14].

The process of drug discovery aims to identify chemical compounds with therapeutic potential for treating human diseases. Despite substantial advances, the success rate for the introduction of new drugs to the market has declined, with the average drug discovery pipeline now exceeding 12 years and costing more than 2 billion dollars [15, 16]. Computational approaches, such as virtual high-throughput screening, are increasingly being used to identify potential lead compounds by simulating and evaluating the binding affinity of numerous compounds against a target [17–20]. Challenges such as the vast combinatorial space of binding poses [21, 22] and ligand conformations [23, 24], coupled with the complex dynamics of these systems [25], limit the effectiveness of traditional virtual screening in reliably producing effective leads. Some computational methods stand out for their efficiency, accuracy, comprehensive assessment of interaction spaces, and broad exploration of chemical space, helping to address the limitations of conventional approaches [26–33]. Although many computational screenings focus on a single protein target, drugs in humans interact with various biological targets through processes such as absorption, distribution, metabolism, and excretion (ADME), which influences their efficacy and safety [31–37]. Considering drug interactions on a proteomic scale could yield more accurate predictions of bioactivity and safety by accounting for both primary and secondary targets, essential for optimizing therapeutic impact and minimizing toxicity.

We developed the Computational Analysis of Novel Drug Repurposing Opportunities (CANDO) platform for multitarget drug discovery, repurposing, and design, aiming to address the limitations of traditional single target, single disease approaches [38–53]. CANDO exploits the fact that drugs approved for human use achieve therapeutic effects and optimal ADME through interactions with multiple biological targets, and that off-target interactions are modulated to minimize adverse drug reactions. CANDO capitalizes on this inherent multitargeting property of small molecules by constructing interaction signatures that reflect drug/compound behaviors across various biological scales. The platform predicts putative drug candidates for every indication/disease by comparing and ranking these interaction signatures in an all-against-all manner, with the hypothesis that drugs/compounds with similar interaction signatures are more likely to display similar biological behavior. The platform is benchmarked by evaluating the recovery of known drug-indication associations in these ranked lists of interaction signatures within specified cutoffs. CANDO therefore deepens our understanding of small molecule therapeutics and their effects on proteins, pathways, and various diseases by leveraging vast multiscale biomedical data on biological systems and the phenotypic impact of their modulation. In addition to rigorous benchmarking [38–53], CANDO and/or its components have been extensively validated prospectively in the context of more than a dozen indications [38, 41, 47, 50–52, 54–64]. Herein, we describe the use of CANDO to predict novel drug candidates for glioma treatment.

## 2 Methods

### 2.1 Applying the CANDO platform for glioma drug discovery overview

We developed a pipeline within the CANDO platform to identify potential drug candidates for glioma (Figure 1). Our approach is based on the hypothesis that drugs/compounds with similar interactions across entire proteomes (“interaction signatures”) are more likely to share therapeutic effects. Signatures were generated by calculating interaction scores between every drug/compound and a comprehensive library of proteins to capture the proteome-wide behaviors of a compound. Compounds with interaction signatures closely matching those of drugs approved for glioma were identified as potential treatments. We benchmarked performance by measuring how frequently known drugs for a given indication were retrieved at various cutoffs in ranked lists of predictions. Next, we compared our glioma specific results against random controls, as well as across all indications. The novel predictions for glioma were then corroborated through literature-based analysis to identify the highest confidence drug candidates. Finally, we conducted a consensus analysis of proteins with the strongest interactions to these novel glioma drug candidates which was further corroborated using protein functional annotations.

**Fig. 1:**
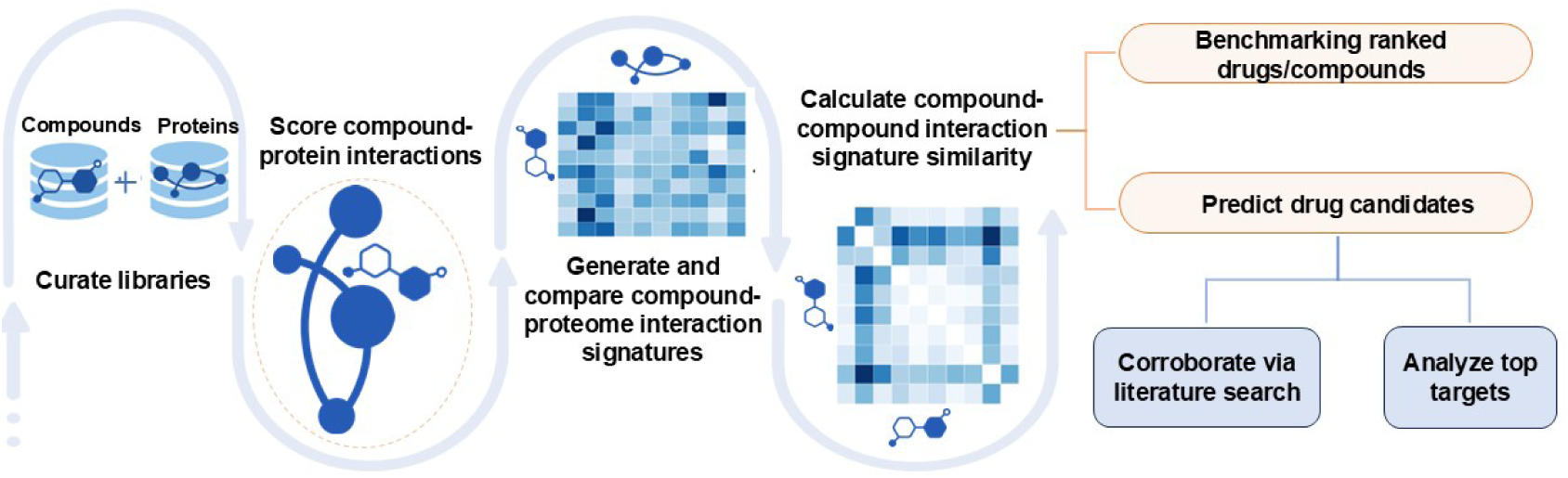
Overview of pipeline for generating novel putative drug candidates for glioma within the CANDO multiscale drug discovery platform. Interaction scores between every protein and drug/compound in the corresponding libraries were calculated using our bioanalytical docking protocol (BANDOCK) [38–53]. This resulted in a compound-proteome interaction signature for each drug/compound describing its functional behavior. Interaction signature similarity was then calculated by comparing pairs of drug-proteome interaction signatures in an all-against-all manner. These interaction signature similarities were sorted and ranked for all drugs approved for an indication and used to benchmark performance and generate putative drug candidates. Benchmarking was conducted by measuring how approved drugs were recovered at various cutoffs. We performed a literature-based analysis to corroborate the glioma drug candidates for their potential to treat this disease. Finally, we identified the protein targets with the strongest interactions to these candidates and further corroborated them using protein functional annotations. The CANDO platform successfully identified multiple candidates demonstrating significant anti-glioma potential, offering a promising avenue to address the current lack of effective treatments for this disease.

### 2.2 Curating compound/protein libraries and indication mapping

Our drug/compound library, sourced mainly from DrugBank [65], comprises 2,449 approved drugs and 10,741 experimental or investigational compounds, totaling 13,457 molecules. The “Homo sapiens AlphaFold2” (AF2) protein library was curated following the application of the AlphaFold2 structure prediction program [66] to the Homo sapiens proteome yielding 20,295 proteins used for this study. The Comparative Toxicogenomics Database (CTD) was used to map the 2,449 approved drugs to 22,771 drug-indication associations based on DrugBank identifiers for drugs and compounds, and Medical Subject Headings (MeSH) terms for approved/associated indications [67, 68]. Benchmarking, which uses a leave-one-out approach (section 2.5), was carried out on indications with at least two approved drugs, yielding a drug-indication mapping consisting of 1,595 indications and 13,226 associations. There were 35 associations in our drug-indication mapping for the indication glioma (MeSH identifier: D005910).

### 2.3 Scoring compound-protein interactions and generating interaction signatures

Interaction scores between each compound and protein were computed using our inhouse bioanalytical docking protocol (BANDOCK); these scores serve as a proxy for binding strength/probability [38, 39, 41, 43, 46, 48]. Binding site prediction was first performed using the COACH algorithm from the I-TASSER suite (version 5.1) [69]. COACH utilizes a library of protein structures bound to ligands, determined through x-ray diffraction, to predict the binding sites and corresponding ligands for each protein based on structural and sequential similarity [70]. BANDOCK then calculates interaction scores by comparing the COACH predicted ligands to the query compound, using similarity between their Extended Connectivity Fingerprints with a diameter of 4 (ECFP4), generated via RDKit [71]. The chemical similarity score is quantified using the Sorenson-Dice coefficient [72], which reflects the similarity between the query compound and the predicted ligand. The highest chemical similarity score is multiplied by the corresponding COACH binding site confidence score to assign an interaction score between a compound and a protein by BANDOCK [38, 39, 41, 43, 46, 48]. BANDOCK is applied between every compound and all proteins in the library, producing compound-proteome interaction signatures describing (compound) behavior.

### 2.4 Calculating ranked compound similarity lists

CANDO calculates all-against-all similarities between compound-proteome interaction signatures to compute drug repurposing accuracy and predict drug candidates [46]. We employed cosine distance for similarity calculations instead of the usual root-meansquare deviation (RMSD) [53] as it enhanced computational speed while maintaining performance. This process was repeated iteratively for all compound pairs in the library, producing a ranked similarity list for each compound.

### 2.5 Benchmarking

Compounds are ranked by the number of times they appear in the similarity lists of the associated drugs above a certain cutoff, resulting in a consensus list. We benchmarked the performance of CANDO by evaluating the recovery of known/approved drugs within similarity lists and aggregated consensus lists across various cutoffs using multiple metrics. The consensus lists classify/rank compounds according to their consensus scores, which reflect how frequently they appear in multiple similarity lists corresponding to all approved drugs for an indication. As mentioned above (section 2.2), we used drug-indication mappings from the Comparative Toxicogenomics Database (CTD) [73] to determine the ranking of approved drugs within specific cutoffs (e.g., top 10, 25, 50, 100) in the similarity and consensus lists of drugs for a given indication with at least two approved drugs [38–53]. Benchmarking performance for all indications, including glioma, was compared to a random control that calculated the probability of correctly selecting an approved drug for an indication by chance, using a hypergeometric distribution [51, 74].

CANDO calculates the following metrics developed in-house to assess performance: indication accuracy, average indication accuracy, new indication accuracy, and new average indication accuracy. Indication accuracy (IA) is the percentage of cases in which at least one approved drug for a given indication appears within a specified rank cutoff in the similarity list of another drug associated with that same indication. Averaging the IA values for all indications with at least two approved drugs produces the average indication accuracy (AIA). New indication accuracy (nIA) captures the frequency with which approved drugs for a given indication appear within particular cutoffs in the consensus list for that indication. The nIA is averaged across all indications to yield the new average indication accuracy (nAIA) metric.

CANDO also calculates the normalized discounted cumulative gain (NDCG) metric, an evaluation measure commonly used in information retrieval to assess the relevance of ranked items based on their positions [75, 76], to evaluate our predictions. In CANDO, NDCG evaluates how effectively a given pipeline prioritizes approved drugs for a specific indication within its similarity lists at specified cutoffs. The NDCG score ranges from 0 to 1, with 1 indicating a perfect ranking [51]. Similarly, the new NDCG (nNDCG) metric assesses the recovery of approved drugs across specified cutoffs in the consensus list for an indication.

### 2.6 Generating drug predictions and corroborating them using literature searches

The CANDO platform was applied to predict novel putative therapeutics for glioma (MeSH identifier: D005910) which had 35 known associations in our drug-indication mapping (section 2.2). As described above, drugs/compounds with interaction signatures similar to those of drugs associated with glioma were ranked. Next, their frequency, or consensus, among the similarity lists was used to identify the top 100 novel drug candidates for glioma. We conducted a literature review using search terms that consisted of the name of each putative drug candidate and “glioma” in Google

Scholar and PubMed. We categorized the candidates as follows: *high-corroboration* for drugs supported by two or more studies showing positive glioma treatment results; *low-corroboration* for drugs targeting glioma-related pathways or supported by a single positive study but lacking confirmation; and *no data found* when no data was present to arrive at any conclusion regarding corroboration.

### 2.7 Analyzing top targets and associated pathways for glioma

We used our in-house top targets protocol to identify the proteins with the strongest interactions with each putative drug candidate that was classified as high-corroboration above. Interaction scores were calculated as described previously (section 2.3) using the BANDOCK protocol, where higher scores (maximum of 1.0) indicate stronger predicted interactions. We then conducted a literature search on Google Scholar and PubMed with the names of the putative drug candidates and proteins to find corroborative evidence supporting the target rationale used by CANDO in generating predictions. We used this information to analyze whether the top targets of the putative drug candidates overlap with proteins in biochemical pathways linked to glioma.

### 2.8 Assessing corroboration between protein functional annotations and predicted top targets

We curated three protein libraries, or “gold standards”, from UniProt [77], GeneCards [78], and a comprehensive literature search to serve as references for evaluating the target predictions for putative glioma treatment candidates. The literature search, data presented in Table 3, focused on identifying targets implicated in glioma from the top targets analysis for high-corroboration putative drug candidates generated by the CANDO platform. The benchmarks assessed the overlap between the gold standard libraries and the top protein targets predicted by CANDO for the top 24 drug candidate predictions. This assessment was repeated with a random set of 24 drug predictions, and the bottom 24 drug predictions as controls. The bottom 24 drug predictions were filtered to include only compounds with at least five heavy atoms to maintain meaningful molecular complexity. To quantify the alignment between the gold standards and the predictions, we employed three key metrics: (A) frequency distribution, (B) percentage overlap, and (C) the Jaccard coefficient, a commonly used metric for assessing similarity across datasets [79, 80]. The Jaccard coefficient calculates the ratio of the intersection and union of two groups and is defined as:

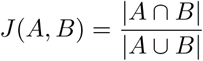

In this context, *A* represents proteins annotated with glioma-related functions, and *B* represents the top protein targets predicted by CANDO. A high Jaccard coefficient indicates that CANDO accurately identifies protein targets that are independently corroborated by functional annotation libraries.

Additionally, we compared the Jaccard coefficient across glioma and other disease indications by using the top predicted targets from the top 24 or top 100 drug candidate predictions for glioma as one group, and functional protein annotations from UniProt as the other group. Selected indications included cancer indications (e.g., metastatic melanoma, non-small cell lung cancer, acute myeloid leukemia) and noncancer diseases (e.g., Alzheimer’s disease, rheumatoid arthritis, asthma). Additionally, we analyzed functional annotations for protein targets in the UniProt database to assess their association with glioma and other disease indications. The Jaccard coefficient was computed separately for glioma-related targets and targets associated with other diseases, including cancer indications (e.g., metastatic melanoma, non-small cell lung cancer, acute myeloid leukemia) and non-cancer diseases (e.g., Alzheimer’s disease, rheumatoid arthritis, asthma). The comparison involved the predicted targets from the top 24 and top 100 drug candidate predictions to evaluate performance differences across indications.

## 3 Results

In summary, the results of this study provided strong evidence for the utility of the CANDO platform in identifying putative drug candidates for glioma. The multitarget approach enabled precise ranking and identification of compounds based on their interaction signatures across the human proteome for treating glioma. The drug candidates exhibited high interaction signature similarity to those of established glioma treatments and were observed to target critical pathways associated with glioma pathogenesis. Benchmarking and comparison with random controls affirmed the robustness of the platform, indicating a high degree of predictive accuracy.

### 3.1 Benchmarking performance

Figure 2 illustrates the benchmarking performance of the CANDO platform for glioma relative to all indications and random controls for both the similarity and consensus lists (section 2.5). The approved drug library returned 35 associated drugs for glioma. Figure 2A shows the AIA and nAIA metrics for all 1,595 indications with at least two approved compounds. AIA ranged from 22% to 44% at the top 10, 25, 50, and 100 cutoffs, outperforming random control accuracies, which ranged from 4% to 26%. CANDO achieved nAIA values ranging from 9% to 26% across the same cutoffs, outperforming the random control for nAIA. The NDCG and nNDCG metrics for all indications, presented in Figure 2B, further validate this performance. NDCG values ranged from 0.044 to 0.059, while nNDCG values varied from 0.049 to 0.083, both exceeding the random control for NDCG.

**Fig. 2:**
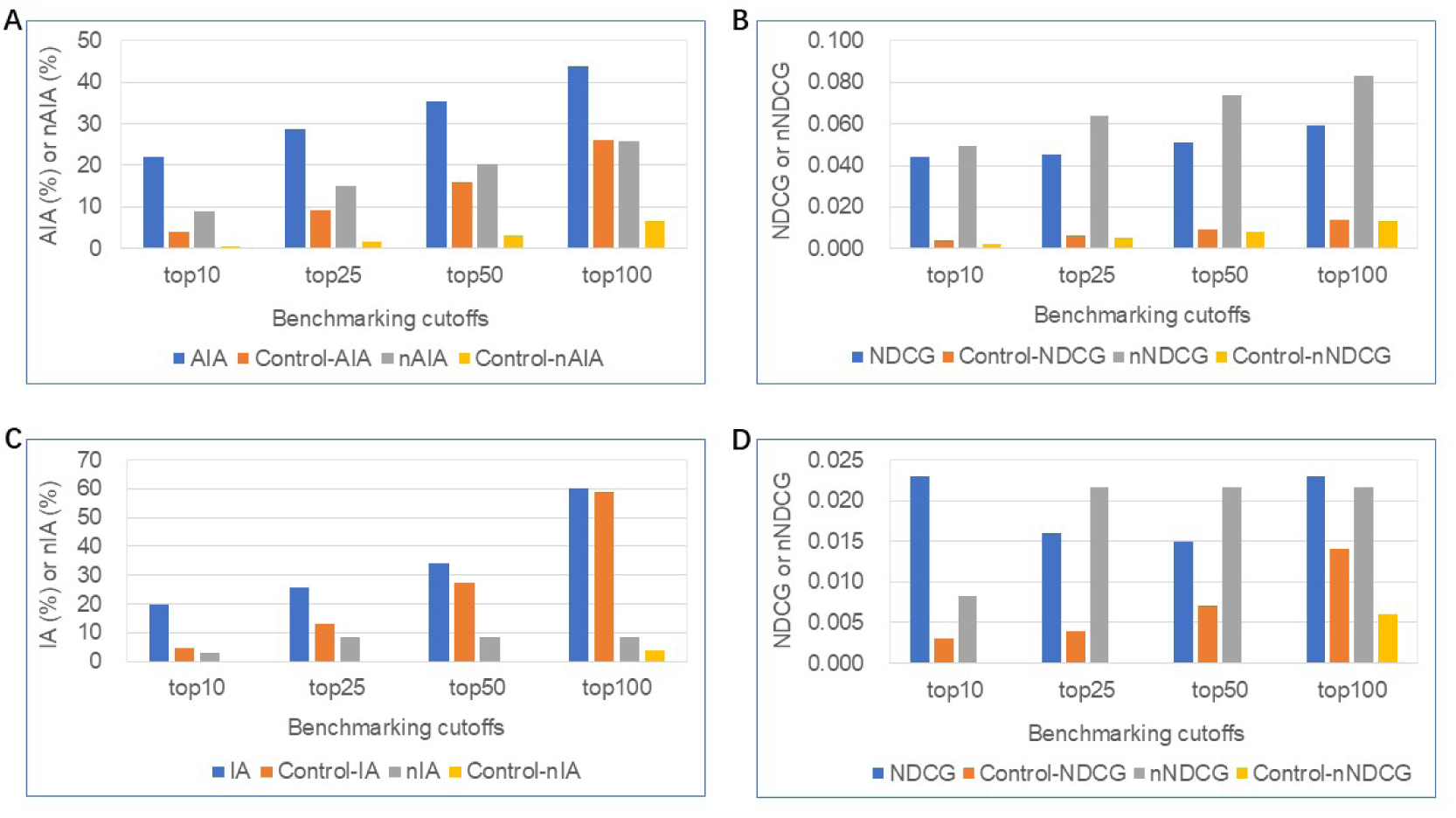
Benchmarking performance of the CANDO platform for glioma relative to all indications and random controls. Performance was evaluated using (A) average indication accuracy (AIA)/new average indication accuracy (nAIA), as well as the normalized discounted cumulative gain (NDCG)/new normalized discounted cumulative gain (nNDCG) metrics across all indications (B). AIA and nAIA at the top 10, 25, 50, and 100 cutoffs, ranging from 22% to 44% and 9% to 26%, respectively, significantly outperform random controls; NDCG and nNDCG metrics, also significantly higher than random controls. In panels C/D, the indication accuracy (IA) and new indication accuracy (nIA) metrics, along with NDCG and nNDCG, were evaluated specifically for glioma. IA ranged from 20% to 60% at the top 10 to top 100 cutoffs, outperforming random controls; NDCG/nNDCG metrics were also higher than random controls. The results indicated that CANDO consistently outperforms random controls in identifying and prioritizing relevant compounds across all indications and glioma-specific predictions.

Figure 2C focuses on glioma-specific benchmarking, where CANDO demonstrated enhanced IA values across all cutoffs compared to controls. The IA for glioma, evaluated using similarity lists, ranged from 20% to 60% across the top 10 to top 100 cutoffs, with a notable top 10 IA of 20%, which is nearly seven times the nIA of 3%. Figure 2D presents the NDCG and nNDCG values for glioma-specific predictions. Glioma has the same NDCG values at the top 10 and top 100 cutoffs, which are both 0.023, while nNDCG values varied from 0.008 to 0.022. In comparison, random controls produced substantially lower NDCG/nNDCG values. The IA/AIA metrics, applied to similarity lists, and the nIA/nAIA metric, specific to consensus lists, collectively demonstrated the robustness of the CANDO platform in leveraging interaction signature similarity and consensus frequency to identify potential drug candidates effectively.

### 3.2 Identifying drug candidates

We used the CANDO platform to predict potential drug candidates for glioma (section 2.6). The 24 most compelling high-corroboration predictions based on ranking metrics from the platform and literature analysis are shown in Table 2. The list of all the top 100 putative drug candidates is given in **Supplementary Materials**. The top ranked drug candidates were Vitamin D compounds: calcifediol, ergocalciferol, and cholecalciferol. Additional drug candidates for glioma included taxanes (cabazitaxel), vinca alkaloids (vinflunine), and topoisomerase inhibitors (topotecan).

**Table 2:**
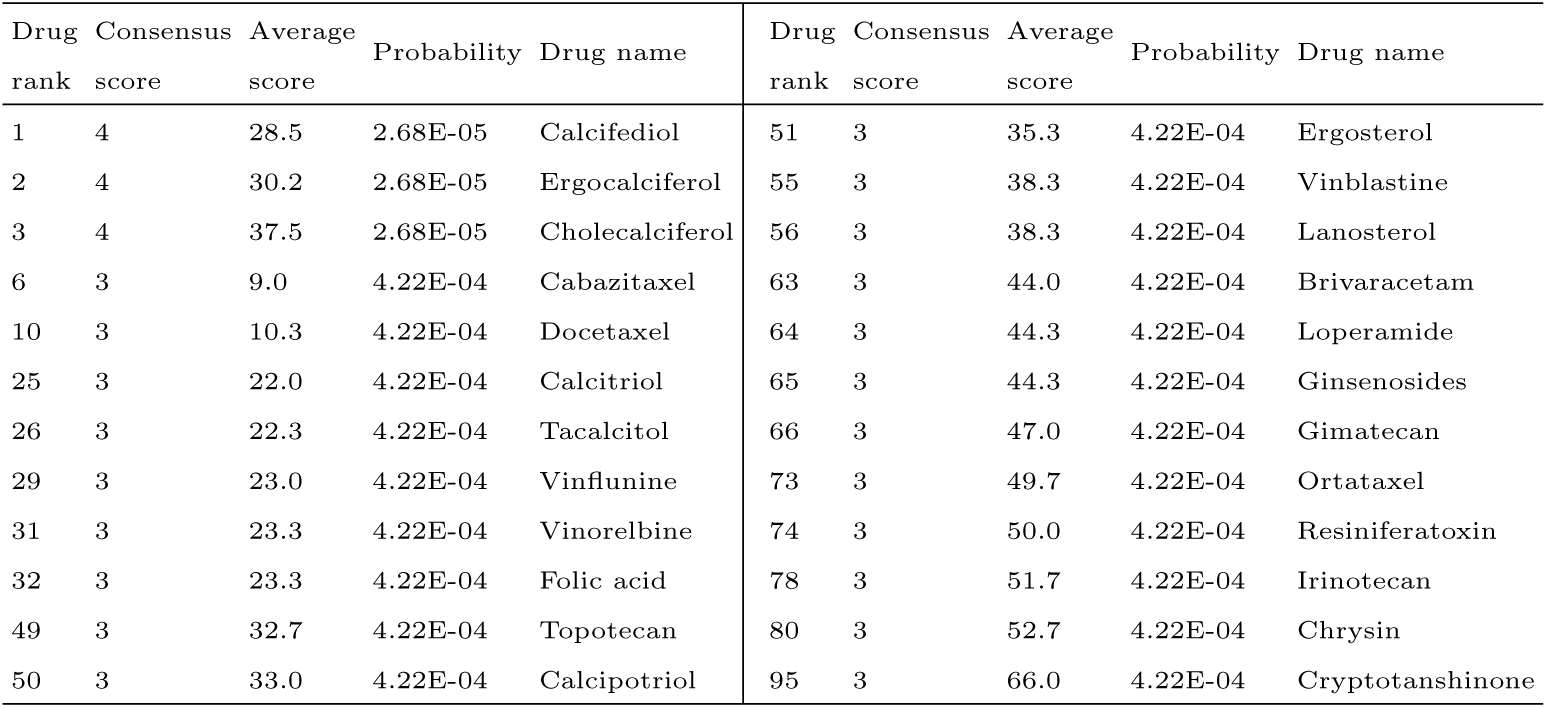
Predicted drug candidates for glioma using CANDO platform that were corroborated using literature analysis. The names of the 24 highcorroboration drug candidates (section 3.2), along with their ranks, consensus/average scores, and probability values are listed. The consensus score represents the number of drug–drug interaction signature similarity lists in which a compound appeared within a particular cutoff. The probability estimates the likelihood of a particular ranked compound appearing by chance, with lower values indicating a better outcome. The overall ranking of a potential drug is determined first by its consensus score and then by its average rank (section 2.6). The best ranked compounds in this consensus list are considered to be the top predictions for an indication. Vitamin D includes a group of compounds such as calcifediol, ergocalciferol, and cholecalciferol, which are ranked as the top three predictions with highest consensus score. This analysis indicates that the signature similarity pipeline within the CANDO platform can generate putative drug candidates for glioma.

| Drug rank | Consensus score | Average score | Probability | Drug name | Drug rank | Consensus score | Average score | Probability | Drug name |
| --- | --- | --- | --- | --- | --- | --- | --- | --- | --- |
| 1 | 4 | 28.5 | 2.68E-05 | Calcifediol | 51 | 3 | 35.3 | 4.22E-04 | Ergosterol |
| 2 | 4 | 30.2 | 2.68E-05 | Ergocalciferol | 55 | 3 | 38.3 | 4.22E-04 | Vinblastine |
| 3 | 4 | 37.5 | 2.68E-05 | Cholecalciferol | 56 | 3 | 38.3 | 4.22E-04 | Lanosterol |
| 6 | 3 | 9.0 | 4.22E-04 | Cabazitaxel | 63 | 3 | 44.0 | 4.22E-04 | Brivaracetam |
| 10 | 3 | 10.3 | 4.22E-04 | Docetaxel | 64 | 3 | 44.3 | 4.22E-04 | Loperamide |
| 25 | 3 | 22.0 | 4.22E-04 | Calcitriol | 65 | 3 | 44.3 | 4.22E-04 | Ginsenosides |
| 26 | 3 | 22.3 | 4.22E-04 | Tacalcitol | 66 | 3 | 47.0 | 4.22E-04 | Gimatecan |
| 29 | 3 | 23.0 | 4.22E-04 | Vinflunine | 73 | 3 | 49.7 | 4.22E-04 | Ortataxel |
| 31 | 3 | 23.3 | 4.22E-04 | Vinorelbine | 74 | 3 | 50.0 | 4.22E-04 | Resiniferatoxin |
| 32 | 3 | 23.3 | 4.22E-04 | Folic acid | 78 | 3 | 51.7 | 4.22E-04 | Irinotecan |
| 49 | 3 | 32.7 | 4.22E-04 | Topotecan | 80 | 3 | 52.7 | 4.22E-04 | Chrysin |
| 50 | 3 | 33.0 | 4.22E-04 | Calcipotriol | 95 | 3 | 66.0 | 4.22E-04 | Cryptotanshinone |

### 3.3 Analyzing targets and pathways related to glioma

The information considered when selecting putative drug candidates for novel treatment included the top (i.e., strongest interaction) protein targets predicted by CANDO, protein and pathway interactions corroborated using the literature, and studies of small molecules in the treatment of glioma observed in the literature 2.7. The top targets predicted by CANDO are outlined in Table 3 and encompass Vitamin D3 receptor, thyroid hormone receptor, acetylcholinesterase, cyclin-dependent kinase 2, tubulin alpha chain, dihydrofolate reductase, and thymidylate synthase. Among these, the strongest interaction was observed between the Vitamin D3 receptor and calcifediol, with a BANDOCK score of 0.850. Figure 3 highlights various important related pathways implicated in the pathogenesis of glioma, including phosphatidylinositol3’-kinase (PI3K)/Akt, mammalian target of rapamycin (mTOR), and Janus kinase (JAK)/signal transducer and activator of transcription (STAT) pathways.

**Fig. 3:**
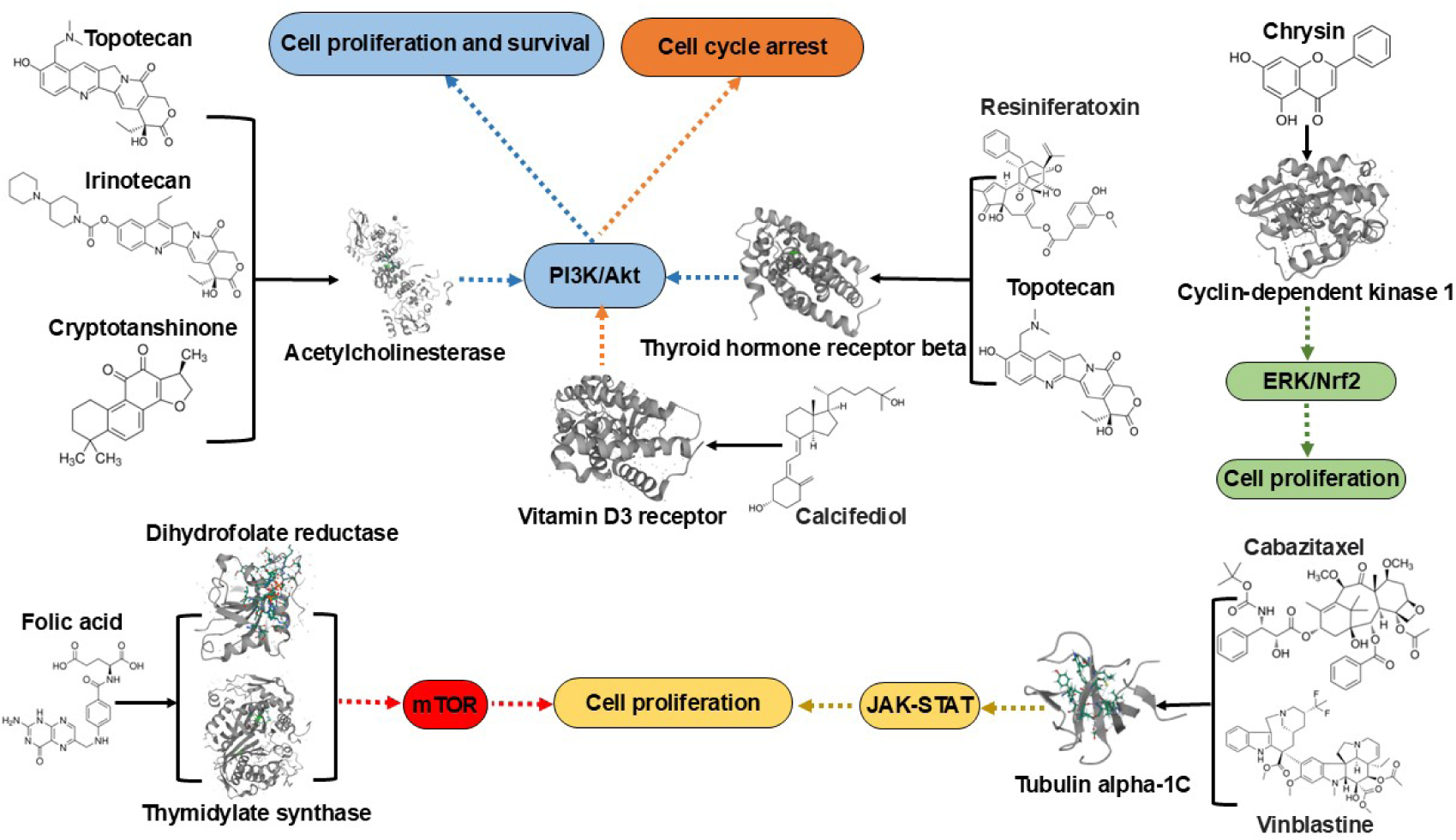
Downstream pathways of top targets for putative drugs for glioma treatment predicted by CANDO. The top targets for putative drugs for glioma are those with the strongest interactions as predicted by our CANDO platform (Table 3) and verified by a functional annotation search (section 3.4). The phosphatidylinositol-3’-kinase (PI3K)/Akt, mammalian target of rapamycin (mTOR), Janus kinase (JAK)/signal transducer and activator of transcription 3 (STAT3) pathways play important roles in the biology of malignant gliomas [81–84]. Topotecan, irinotecan, and cryptotanshinone all interact with acetylcholinesterase (ACHE), ranking 8th (topotecan and irinotecan) and 1st (cryptotanshinone), respectively. Compounds resiniferatoxin and topotecan strongly interact with the thyroid hormone receptor beta (THRB). The targets ACHE and THRB both influence glioma cell proliferation and survival by regulating the PI3K/Akt signaling pathway (blue). Cell cycle arrest is one of the most well-studied mechanisms accounting for the antitumor activity of vitamin D in gliomas (orange). The compound chrysin interacts with cyclin-dependent kinase 1 (CDK1), targeting glioma cell proliferation via the ERK/Nrf2 signaling pathway (green). Dihydrofolate reductase (DHFR) and thymidylate synthase (TYMS) are key targets of folic acid, modulating glioma cell proliferation through the mTOR signaling pathway (red). Taxanes (e.g., cabazitaxel) and vinca alkaloids (e.g., vinblastine) interact with tubulin alpha-1C, influencing glioma through the JAK-STAT pathway (yellow). Our study allows for comprehensive mechanistic understanding of drug candidate behavior across multiple scales, showcasing the CANDO platform’s capability to accurately identify novel drug candidates and their mechanisms through a multifaceted strategy.

### 3.4 Determining overlap between protein functional annotations and CANDO predicted top targets

Figure 4A illustrates the frequency distribution of overlaps between our three gold standard protein libraries and the top protein targets predicted by CANDO. For all gold standards, targets of the top 24 drug candidates showed the highest proportion of overlaps within the highest ranked bin (1-20). In contrast, targets from the random 24 drug candidates and bottom 24 drug candidates exhibited a comparatively uniform distribution across the 5 bins. Figure 4B presents the cumulative percentage overlap as a function of rank cutoff for the predicted targets across the gold standard libraries. The targets from the top 24 drug candidate predictions demonstrated a near-saturation of overlap at lower rank cutoffs (e.g., 80% overlap by rank 20 for Table 3 and UniProt), emphasizing their strong alignment with gold standard targets. In contrast, the targets from the random 24 and bottom 24 drug candidate predictions exhibited a slower increase in overlap percentage, with cumulative overlaps remaining below 10% even at a rank cutoff of 100 for Table 2 and GeneCards. As shown in Figure 4C, the Jaccard coefficient values further corroborate the findings from the frequency distribution and overlap percentage analyses. Across all libraries, the Jaccard coefficient for the top protein targets from the top 24 drug candidate predictions was consistently higher compared to those derived from the random 24 or bottom 24 drug candidate predictions.

**Fig. 4:**
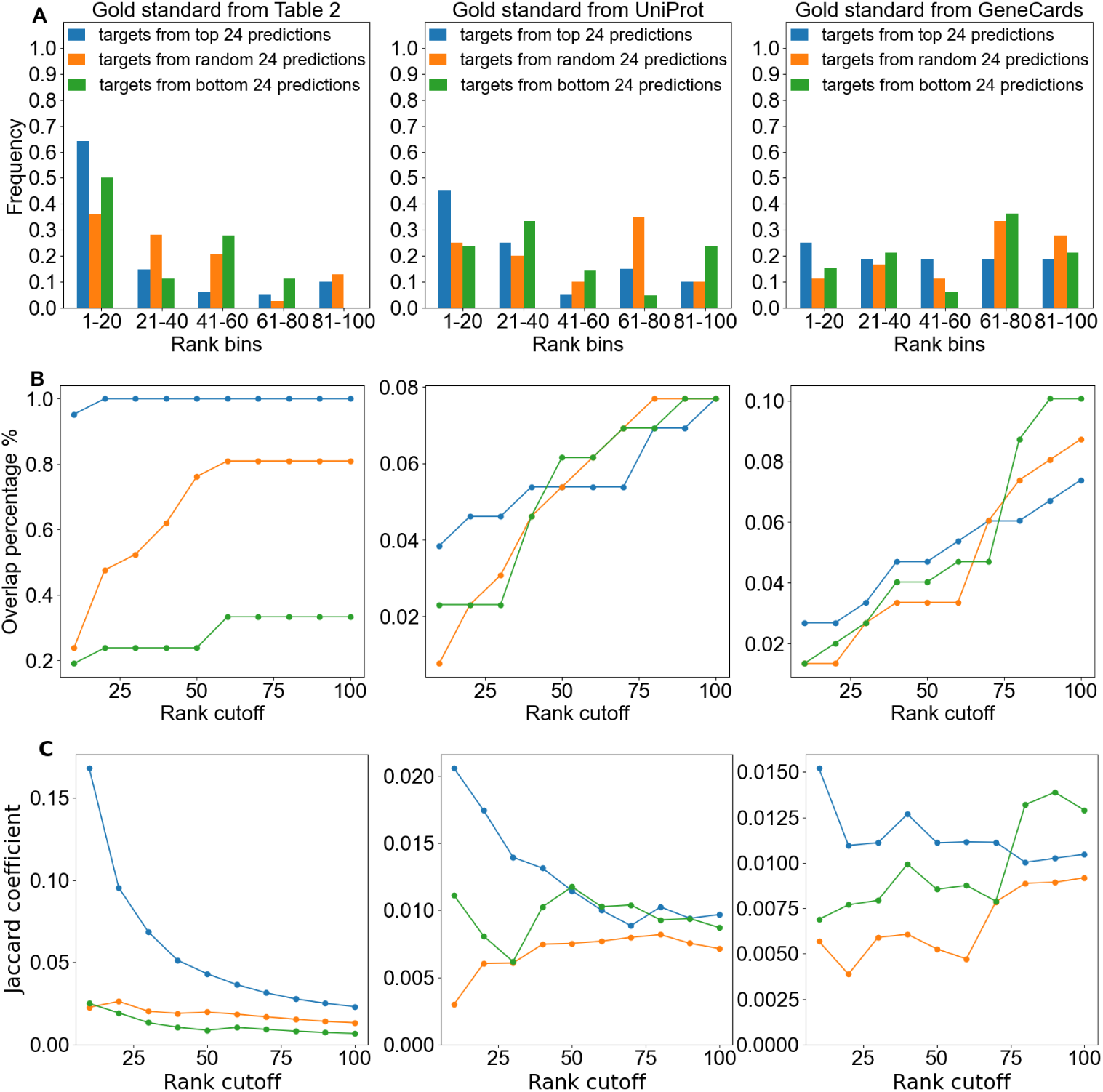
Overlap between protein functional annotations and CANDO predicted top targets across gold standard libraries. This figure compares the overlap between CANDO predicted targets and gold standard annotations (Table 3, UniProt, and GeneCards) for glioma-related proteins. A) Frequency distributions show the proportion of predicted targets that overlap with gold standard proteins within rank bins for top 24 drug candidate predictions (blue), random 24 drug candidate predictions (orange), and bottom 24 drug candidate predictions (green). B) The line graphs show the percentage of gold standard proteins that overlap with prediction targets as a function of rank cutoff (from 10 to 100). C) Targets from the top 24 drug candidate predictions demonstrate a higher Jaccard coefficient compared to from random 24 and bottom 24 drug candidate predictions across all gold standards. The Jaccard coefficient quantifies the similarity between CANDO predicted targets and gold standard targets. Each column corresponds to a different gold standard: Table 3 (left), UniProt (center), and GeneCards (right). Results demonstrate targets from the top 24 drug candidate predictions generally reflect a stronger signal compared to that of random or bottom drug candidate predictions, highlighting the predictive accuracy of the top candidates.

We found that the Jaccard coefficient for top (rank *≤* 10) predicted targets of the top 24 and top 100 drug candidate predictions for glioma was higher when compared to UniProt glioma protein functional annotations (Figure 5). In contrast, the Jaccard coefficient was lower when comparing glioma targets to protein functional annotations for other indications. Indications demonstrating a lower Jaccard coefficient include other cancer indications such as non-small cell lung cancer and metastatic melanoma, as well as non-cancer diseases like Alzheimer’s disease and rheumatoid arthritis. This result suggests that the top predicted glioma targets identified by CANDO are more functionally relevant to glioma-related gold standard protein targets than those of other indications, highlighting the effectiveness of the pipeline in identifying meaningful targets. When compared to the broader rank distribution shown in the earlier line plot (Figure 4), the rank *≤* 10 results highlight the ability of the pipeline to capture high-confidence and/or known targets for glioma. This trend underscores the utility of using stringent rank cutoffs to identify highly specific target overlaps, particularly for glioma.

**Fig. 5:**
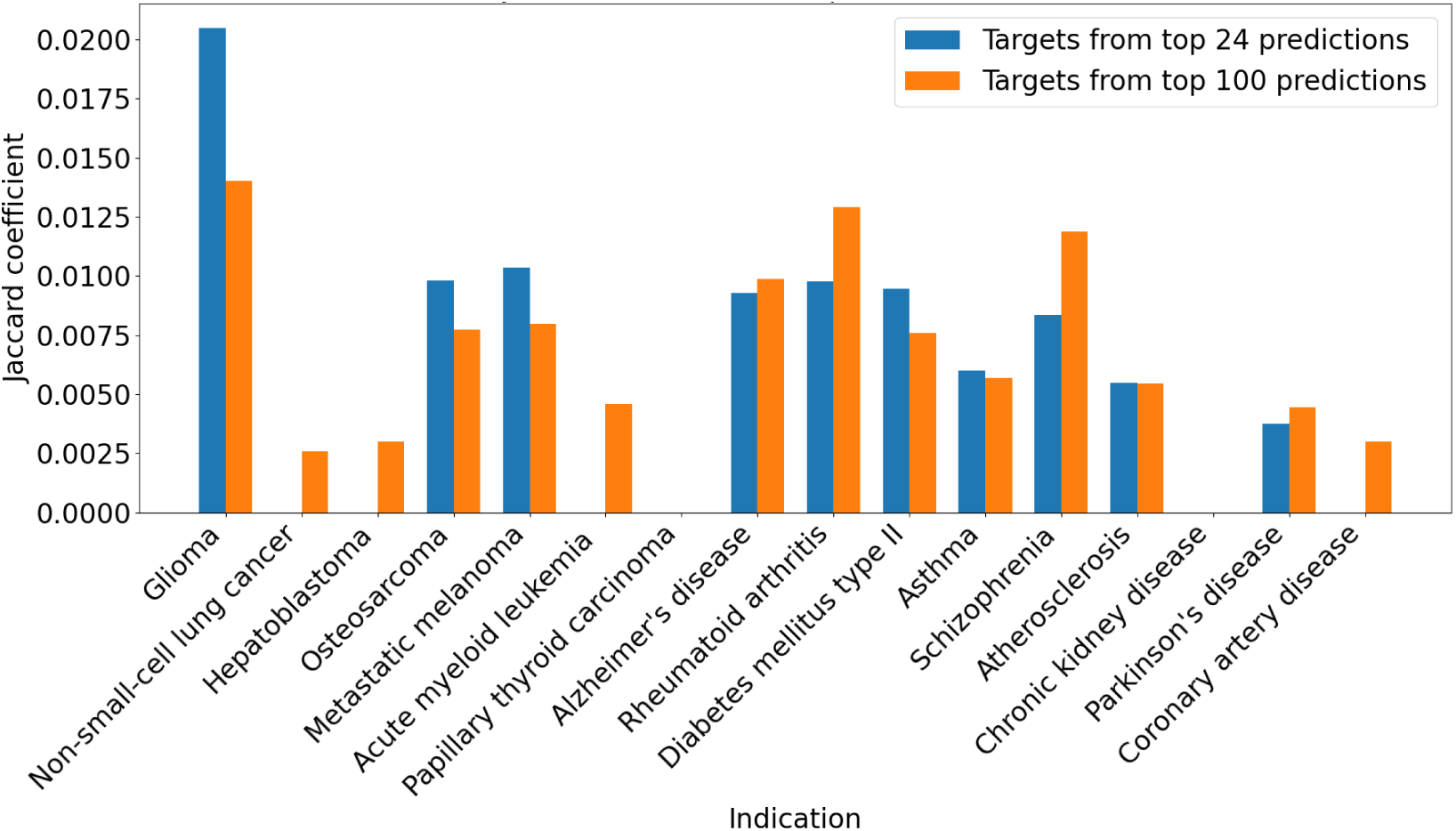
Overlap between protein functional annotations and CANDO predicted top targets across indications. This bar chart compares the overlap between targets with the strongest predicted interactions to our top drug candidates and existing protein annotations across glioma and other indications, using the Jaccard coefficient (vertical axes). The Jaccard coefficient quantifies the overlap between protein functional annotations (from UniProt) and CANDO-predicted drug targets. Two comparisons are made: the overlap with top 24 drug candidate predictions (blue) and the overlap with top 100 drug candidate predictions (orange). A higher coefficient indicates stronger alignment between the predicted and known targets. The horizontal axis lists the various indications, including cancers (glioma, hepatoblastoma, metastatic melanoma) and non-cancer conditions (diabetes mellitus type II, coronary artery disease, and chronic kidney disease). Results show that the Jaccard coefficient for glioma is notably higher than that of other indications, highlighting the effectiveness of the CANDO platform in identifying glioma-related protein targets.

**Table 3:**
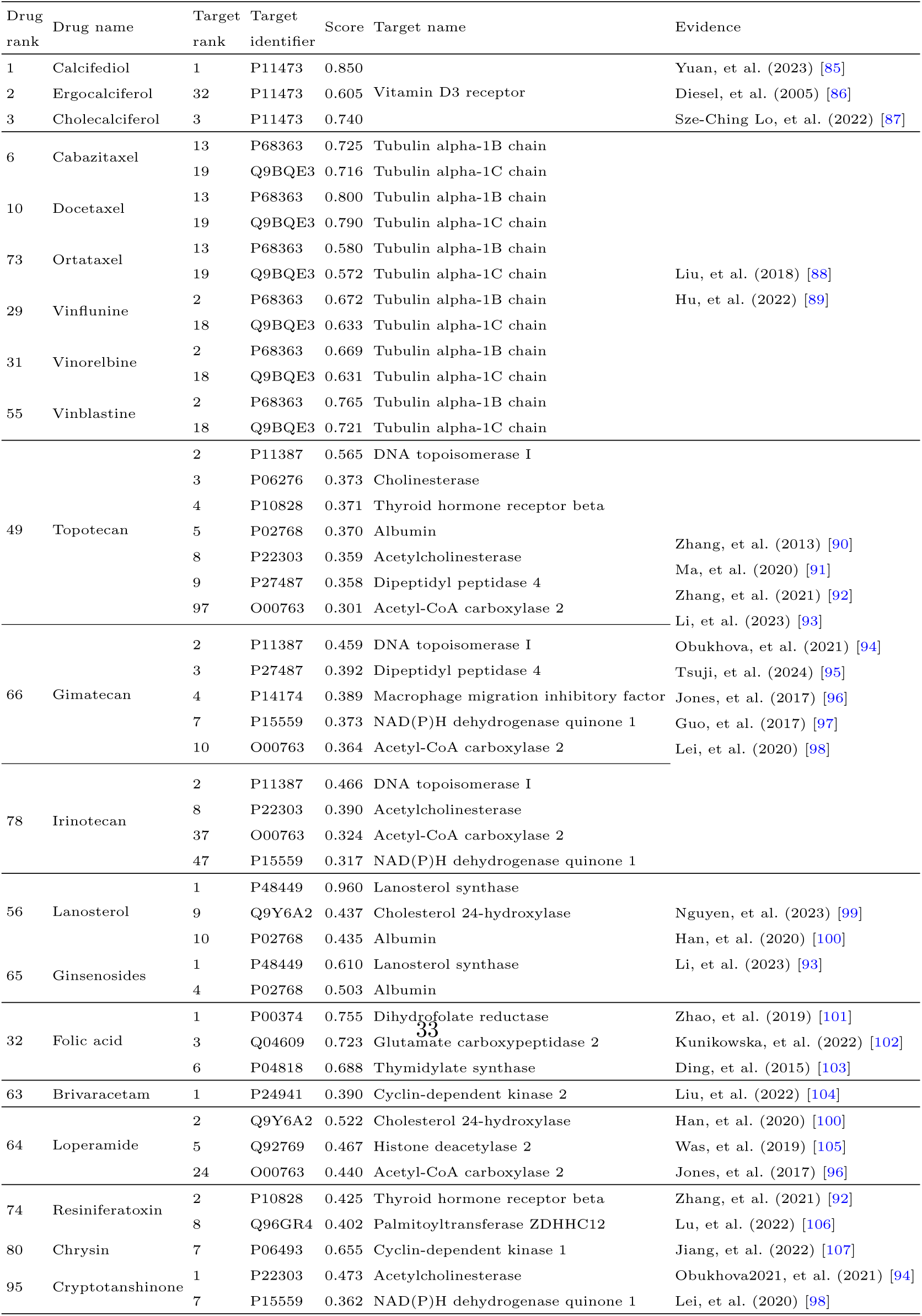
Top targets analysis for high-corroboration putative drug candidates for glioma generated by the CANDO platform. The name of the high-corroboration drug candidates, their predicted ranks based on consensus scoring (section 2.6), the rank of the target from the top targets analysis (section 2.7), the target UniProt identifier, the predicted interaction score between these predictions and targets, the target name, and the evidence we found for the target being implicated in glioma, are listed. Higher scores indicate a higher likelihood of interaction. From this analysis, we highlighted Vitamin D3 receptor, thyroid hormone receptor, acetylcholinesterase, cyclin-dependent kinase 1, tubulin alpha chain, dihydrofolate reductase and thymidylate synthase as the most promising targets for glioma.

## 4 Discussion

CANDO identified potential glioma treatments that included drugs approved for other indications such as vitamin D (calcifediol), taxanes (cabazitaxel and docetaxel), vinca alkaloids (vinblastine and vinflunine), topoisomerase inhibitors (topotecan and irinotecan), and folic acid. Additionally, investigational compounds like ginsenosides, brivaracetam, chrysin, resiniferatoxin, and cryptotanshinone were also identified as promising drug candidates (Table 2). Literature-based analysis was conducted to corroborate these potential drugs and compounds for glioma, examining supporting evidence for their targets and pathways (Table 3 and Figure 3). The top drug candidates generated via the interactomic signature pipeline of CANDO may be exerting their therapeutic effects by impacting multiple pathways implicated in glioma. We examined the top drugs/compounds and targets predicted by CANDO in further detail, comparing and contrasting to what is known about their relevance to glioma in the literature; a detailed description follows below.

### 4.1 Vitamin D3 receptor metabolites

Vitamins may have a role in the etiopathogenesis of central nervous system (CNS) cancers [108]. Vitamin D comprises a group of fat-soluble steroids, with vitamin D3 (cholecalciferol) and vitamin D2 (ergocalciferol) being the most significant [109]. Calcifediol (25-hydroxyvitamin D3),is the precursor for calcitriol, the active form of vitamin D [110]. Recent research suggests that the levels of the progenitor of calcitriol correlate with progression of glioma [111–114]. Cholecalciferol has shown promise in glioma treatment, especially glioblastoma multiforme (GBM), due to its ability to regulate cell cycle biomarkers and enhance the anti-tumor immune response [85, 115]. Studies indicate that vitamin D analogs, including ergocalciferol, could modulate biomarkers involved in cell cycle regulation and apoptosis in glioblastoma [115]. Cell cycle arrest is one of the most well-studied mechanisms accounting for the anti-tumor activity of vitamin D in gliomas. Vitamin D has been shown to induce antiglioma effects through cell cycle arrest and the phosphoinositide 3-kinase (PI3K)/Akt pathway [87].

### 4.2 Taxanes

Taxanes are a class of diterpenes commonly used as chemotherapy agents, mainly including cabazitaxel, docetaxel and paclitaxel [116–118]. Cabazitaxel is a secondgeneration semisynthetic taxane. Contrary to other taxane compounds, cabazitaxel is a poor substrate for P-gp efflux pump; therefore, it easily penetrates across the BBB [119, 120]. Cabazitaxel shows a significant inhibitory effect on glioma [121, 122]. Other studies have reported that cabazitaxel exerts its anti-proliferative effects on cancer cells by binding to tubulin [123]. One study indicates that tubulin alpha-1C chain (TUBA1C) may potentially regulate the pathogenesis of glioma through Janus kinase (JAK)/signal transducer and activator of transcription (STAT) (JAK-STAT) pathway [124]. Docetaxel, a taxane-class anti-mitotic agent, demonstrates the ability to induce cell apoptosis in glioma and shows substantial inhibitory activity against tumor growth [125]. Furthermore, it is recognized as one of the leading drug candidates for brain tumor therapy [126]. In our study, both cabazitaxel and docetaxel are predicted to strongly interact with TUBA1C, with predicted interaction scores of 0.716 and 0.790, respectively (Table 3).

### 4.3 Vinca alkaloids

Vinca alkaloids are a class of chemotherapy agents with anti-mitotic and antimicrotubule properties, including compounds such as vinflunine, vinorelbine, vinblastine, and vincristine [127–129]. Vinflunine, a fluorinated vinca alkaloid, disrupts microtubule dynamics, a process essential for cell division, and has shown potential for glioma treatment [130, 131]. Vinorelbine, a semi-synthetic vinca alkaloid, is an anti-mitotic chemotherapy drug used to treat various cancers, including breast cancer, non-small cell lung cancer, and glioma [132]. Its antitumor effect arises from its ability to inhibit mitosis by interacting with tubulin [133]. In 2000, a pilot study of weekly vinblastine in patients with recurrent low-grade gliomas (LGG) yielded promising results [134, 135]. Compared to vinflunine and vinorelbine, vinblastine demonstrated a higher interaction score with the TUBA1C target (Table 3).

### 4.4 Topoisomerase I inhibitors

Topoisomerase inhibitors are chemical compounds that block the action of topoisomerases, which are broken into two broad subtypes: type I topoisomerases (TopI) and type II topoisomerases (TopII) [136, 137]. TopI inhibitors, like topotecan, are watersoluble camptothecin analogs that have shown cytotoxicity toward a variety of tumor types [138]. Topetecan can pass through the BBB and exhibits significant activity in treating glioblastoma multiforme[139, 140]. Additionally, it has been observed to induce cell cycle arrest at the G0/G1 and S phases [90, 141]. Irinotecan (CPT-11), a prodrug of 7-Ethyl-10-hydroxycamptothecin (SN-38), shows some antitumor activity in patients with recurrent glioblastoma multiforme, with response rates of 0 to 17% in several trials [142, 143]. Gimatecan is a lipophilic oral camptothecin analogue with preclinical activity in glioma models [144].

### 4.5 Folic acid

Folic acid (FA) targets the folate receptor (FR), which is overexpressed on the cell surface of various cancer cells [145, 146]. Folate supplementation, particularly at high doses, has been suggested to have cytotoxic effects on glioma cells, making it a potential candidate for further exploration in glioma therapies [147]. In addition, utilizing lidocaine liposomes modified with folic acid has been demonstrated to suppress the proliferation and motility of glioma cells [148]. One clinical research study explored the role of folate-related compounds, such as L-methylfolate, in combination therapies for glioma, showing potential epigenetic modifications and enhanced sensitivity to standard treatments like temozolomide [149]. Zhao, et al. [101] hypothesized that inhibition of dihydrofolate reductase/thymidylate synthase might modulate the cell sensitivity of glioma cells to temozolomide through the mTOR signaling pathway. DHFR and TYMS are key metabolic enzymes in the folic acid signaling pathway, with high predicted interaction scores of 0.755 and 0.688, respectively, to folic acid in this study (Table 3).

### 4.6 Other drug candidates and key target interactions

Ginsenosides, active components found in Panax ginseng, show potential in glioma treatment due to their various therapeutic properties, including anticancer and neuroprotective effects [150, 151]. Additionally, ginsenoside has been shown to inhibit the growth of human glioma U251 cells, promoting apoptosis and affecting key signaling pathways involved in cell survival and death [152]. Additionally, compounds such as brivaracetam, which lack enzyme-inducing activity on the cytochrome system, could be considered promising candidates for addressing brain tumor-related epilepsy [153]. Chrysin, an active natural bioflavonoid, is predicted to target cyclin-dependent kinase 1 with an interaction score of 0.655 (Table 3), and has been proven to protect against carcinogenesis [107]. Cyclin-dependent kinase is the target for glioma cell cycle arrest at G2 and M phases [154]. Chrysin exerts anticancer activity in glioblastoma cell lines possibly via the ERK/Nrf2 signaling pathway [155]. Resiniferatoxin, a naturally occurring irritant tricyclic diterpene which combines structural features of the phorbol ester tumor promoters and of capsaicin [156]. It activates transient vanilloid receptor (TRP), which was previously associated primarily with cardiovascular and neuronal regulation, but might also present avenues for exploration in glioma pathogenesis [157]. We observed evidence of interaction between resiniferatoxin and the thyroid hormone receptor beta target (Table 3 and Figure 3). Cryptotanshinone is one of the main representative components isolated from the roots of Salvia miltiorrhiza. Lu, et al. [158] indicated that cryptotanshinone can inhibit human glioma cell proliferation.

As shown in Table 3 and Figure 3, we predicted interactions between topotecan, irinotecan, and cryptotanshinone and acetylcholinesterase (AChE), a newly recognized marker for glioma (Table 3). One study reported that irinotecan or its metabolites directly interact with AChE, inhibiting the conversion of acetylcholine to choline, which leads to an accumulation of acetylcholine and subsequent cholinergic syndrome symptoms [159]. Bioinformatic analysis has shown that AChE is connected to proteins in the PI3K/Akt pathway, which promotes anti-apoptotic and proliferative effects in brain tumors [94]. There is limited evidence of interactions between topotecan, irinotecan, cryptotanshinone, and AChE; our study therefore provides predictive evidence of these interactions. Additionally, predictions of topotecan and resiniferatoxin targeting thyroid hormone receptor beta are novel to this study. The thyroid hormone receptor influences glioma progression by regulating the PI3K/Akt signaling pathway [92]. Therefore, candidates targeting the PI3K/Akt pathway may hold promise for glioma treatment.

Our protein list, derived from predictions, highlights glioma-relevant targets but is inherently incomplete, similar to databases like UniProt or GeneCards, as each captures only a partial view of glioma biology. While this list serves as one of the curated gold standards for our analysis, incorporating known treatment targets in future studies could provide a more comprehensive benchmark. Limitations of this study include the arbitrary rank cutoffs, which may exclude moderately ranked targets that overlap meaningfully with gold standard libraries, and the use of the Jaccard coefficient, a binary metric that overlooks relative ranks or prediction scores. Additionally, our focus on glioma leaves the robustness of this approach across other indications underexplored, particularly for diseases with fewer validated targets. Finally, the analyses may bias toward frequently predicted top targets, underrepresenting less common targets with potential therapeutic value. To address these limitations, future studies will integrate score-based cutoffs, and consider a broader range of rank and score distributions.

## 5 Conclusions

We utilized our CANDO platform to explore potential novel treatments and their associated protein targets for glioma. By integrating a combination of computationally generated and experimentally observed data from benchmarking, prediction, corroboration of putative drug candidates using literature-based searches, top protein target analysis, and protein functional annotation, we identified promising treatments for glioma, including Vitamin D, taxanes, vinca alkaloids and topoisomerase inhibitors. Additionally, we highlighted several protein targets and related pathways linked to glioma, including Vitamin D3 receptor, thyroid hormone receptor, acetylcholinesterase, cyclin-dependent kinase 2, tubulin alpha chain, dihydrofolate reductase and thymidylate synthase. This study offers insights into the potential mechanisms underlying glioma and demonstrates the potential of the CANDO platform in identifying effective treatments against this disease.

## Acknowledgements

The authors would like to acknowledge the Center for Computational Research (CCR) at University at Buffalo for computational support. We would also like to thank all members of the Samudrala Computational Biology Group.

## Declarations

- Funding: This work was supported in part by a National Institutes of Health (NIH) Director’s Pioneer Award (DP1OD006779), a NIH Clinical and Translational Sciences (NCATS) Award (UL1TR001412), a NIH National Library of Medicine (NLM) T15 Award (T15LM012495), a NIH NLM R25 Award (R25LM014213), a NIH NCATS ASPIRE Design Challenge Award, a NIH NCATS ASPIRE Reduction-to-Practice Award, a National Institute of Standards of Technology (NIST) Award (60NANB22D168), a NIDA Mentored Research Scientist Development Award (K01DA056690), and startup funds from the Department of Biomedical Informatics at the University at Buffalo.
- Competing interests: The funders had no role in the design of the study; in the collection, analyses, or interpretation of data; in the writing of the manuscript, or in the decision to publish the results. The authors have formed multiple startups that seek to commercialise the outputs of the CANDO platform.
- Ethics approval and consent to participate: Not applicable
- Consent for publication: Not applicable
- Data availability: Data including compound-protein interaction matrices are available upon request.
- Materials availability: Not applicable
- Code availability: Not applicable
- Author contribution: SM.X. helped conceive the research project and pipelines, conducted all analyses, and drafted the manuscript. W.M., A.E., M.V.N., and YK.H. helped data analyses. Z.F. and R.S. conceived of the research design, approach, and methods, supervised the overall project, and edited the manuscript. All authors have read and agreed to the published version of the manuscript.

## List of Abbreviations

CANDO: Computational Analysis of Novel Drug Opportunities
BBB: Blood-brain barrier
P-gp: P-glycoprotein
ADME: Absorption, distribution, metabolism, and excretion
BANDOCK: Bioanalytical docking protocol
AF2: AlphaFold2
CTD: Comparative Toxicogenomics Database
MeSH: Medical Subject Headings
ECFP4: Extended Connectivity Fingerprints with a diameter of 4
RMSD: Root-mean-square deviation
IA: Indication accuracy
AIA: Average indication accuracy
nIA: New indication accuracy
NDCG: Normalized discounted cumulative gain
nNDCG: New NDCG
PI3K: Phosphatidylinositol-3’-kinase
mTOR: Mammalian target of rapamycin
JAK: Janus kinase
STAT: Signal transducer and activator of transcription
STAT3: Signal transducer and activator of transcription 3
AChE: Acetylcholinesterase
THRB: Thyroid hormone receptor beta
CDK1: Cyclin-dependent kinase 1
DHFR: Dihydrofolate reductase
TYMS: Thymidylate synthase
CNS: Central nervous system
GBM: Glioblastoma multiforme
TUBA1C: Tubulin alpha-1C chain
LGG: Low-grade gliomas
TOPI: Type I topoisomerases
TOPII: Type II topoisomerases
CPT-11: Irinotecan
SN-38: 7-Ethyl-10-hydroxycamptothecin FA Folic acid
FR: Folate receptor
TRP: Transient vanilloid receptor
NIH: National Institutes of Health
NCATS: NIH Clinical and Translational Sciences
NLM: NIH National Library of Medicine
NIST: National Institute of Standards of Technology
CCR: Center for Computational Research

